# Expanding the gain-variance Pareto via optimal recycling and genomic mating

**DOI:** 10.1101/2025.09.28.679020

**Authors:** Seifelden M. Metwally, Javier Fernández-González, Julio Isidro y Sánchez

## Abstract

The optimization of mating plans, or optimal genomic mating (OGM), is a powerful breeding strategy that balances genetic gain with the preservation of diversity, securing long-term improvement. However, existing OGM implementations neglect the recycling stage, where naïve truncation selection dissipates the diversity initially safeguarded. Here, we propose an integrated strategy that couples optimal recycling with genomic mating to better control genetic diversity while delivering competitive genetic gains. Using stochastic simulations of line and hybrid breeding schemes, we show that the integrated strategy retained 1.6–2.0 times more diversity than OGM alone and 3.5–5.0 times more than truncation mating based on family means and the usefulness criterion (UC). These were equivalent to maintaining around 1.7 and 2.4 times less realized inbreeding rates. Additionally, it improved the efficiency of translating variance into gain by 21.6–49.8% and 67.4–108.2% compared to the sole implementation of OGM and truncation mating strategies. We also demonstrate the utility of our newly developed intuitive and standardized metric, *proportion of additive standard deviation lost* (PropSD), for managing diversity in the crossing and recycling stages. Pareto optimal solutions were achieved at around 2–3% and 4–5% PropSD without and with optimal recycling. Finally, we derive a closed-form expression quantifying the expected advantage of UC over mean-based mating. Modeling within-family variance offered limited additional benefit, mainly due to high family mean-to-standard-deviation variance ratios. Overall, our proposed framework advances genomic selection programs to be sustainable by effectively preserving genetic diversity for future genetic improvement.

## Introduction

Genetic diversity has long been framed in terms of inbreeding and regarded as the entity that must be sacrificed in the pursuit of gain (Lindgren and Mullin 1997). Early work sought to reduce inbreeding rates without compromising genetic progress (Wray and Thompson 1990; Toro and Perez-Enciso 1990; Wray and Goddard 1994; Caballero et al. 1996; Meuwissen 1997; Grundy et al. 1998). The advent of genomic selection (GS) intensified this challenge, as it increases selection intensity (*i*), improves accuracy (*ρ*), and shortens generation intervals (*L*) (Alemu *et al*. 2024). However, these advantages also accelerate allele fixation, since selection tends to favor individuals closely related to the elite members of the training set. Over time, this erodes the reservoir of “dark diversity” needed for future breeding gains (Jannink 2010; De Beukelaer et al. 2017).

One solution is to control mating in the crossing block stage of breeding programs. Optimal contribution theory selects parents based on their expected genetic contributions, subject to predefined pedigree-based constraints on group coancestry (Meuwissen 1997). Building on this, Kinghorn (2011) optimized contributions and mating simultaneously (optimal mate allocation, OMA) using a differential evolution algorithm (Storn and Price 1997), to account for practical constraints that could hinder the realization of optimal contributions and leverage specific combining abilities (Toro and Varona 2010; Endelman 2025). Genomic relationship matrices (*G*) soon replaced pedigree-based (*A*) matrices (Woolliams *et al*. 2015), and diverse OMA variants emerged. Akdemir and Sánchez (2016) coined the term *genomic mating* and used *G* to estimate breeding values, progeny variance, and group coancestry for optimization. `AlphaMate` (Gorjanc and Hickey 2018) extended it to accommodate genome editing, while `SimpleMating` (Peixoto *et al*. 2024a) enabled multitrait optimization with the usefulness criterion (UC) (Schnell and Utz 1975). Later, Endelman (2025) introduced convex OMA (`COMA`) to extend the functionality of OMA to polyploid species. Most recently, the `MateR` package (Fernandez-Gonzalez *et al*. 2025) was developed to unify prediction of family mean and variance across ploidy levels under both *genotypic* and *breeding* parameterizations (Vitezica *et al*. 2013), while also introducing an interpretable diversity metric directly linked to additive variation and long-term gain, the *proportion of additive standard deviation lost* (PropSD).

Despite these advances, the recycling stage remains under-optimized. It is plausible to expect that a considerable amount of diversity saved in the crossing block is lost by truncation recycling, which favors highly related top performers. Instead, *optimal recycling* should be performed to recycle individuals that balance gain and variance. Here, we present an integrated strategy, implemented in `MateR` (Fernandez-Gonzalez *et al*. 2025) and powered by TrainSel genetic and simulated annealing algorithms (Akdemir *et al*. 2021), which optimizes mating and recycling to better control genetic diversity (Figure 1).

**Figure 1.**
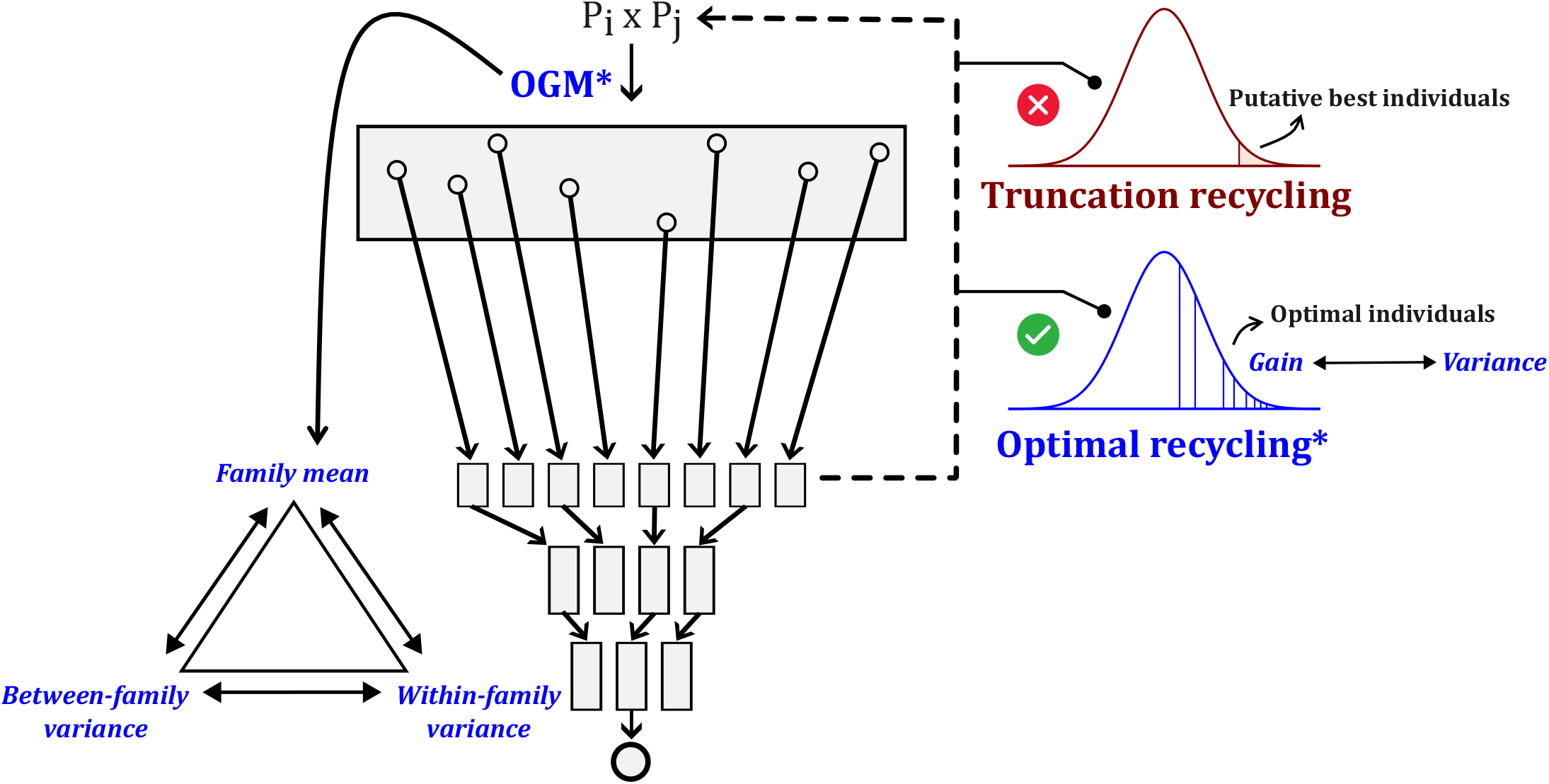
Graphical abstract showing a generic genomic selection breeding scheme (modified from Bančič *et al*. (2025)) and the concept of our integrated optimal recycling and genomic mating strategy. Steps marked with an asterisk (*) denote processes executed by the `MateR` package. **OGM**: optimal genomic mating at the crossing block stage.

This study has three main objectives: (i) we test whether combining optimal recycling with genomic mating outperforms genomic mating alone; (ii) demonstrate the use of the PropSD to control diversity in different breeding systems and program stages (crossing and recycling); (iii) assess the added value of modeling within-family variance in optimal genomic mating (OGM), and provide a mathematical derivation to quantify the importance of UC.

## Theory

In this section, we describe how the family mean and standard deviation are predicted by `MateR` in this study, together with the derivation of PropSD and the quantification of the importance of UC. Throughout, we adopt the *genotypic* parameterization of additive and dominance effects as described by Vitezica *et al*. (2013).

### Prediction of family mean

We compute the family mean as the expected genotypic value obtained by averaging single-locus contributions with respect to genotype frequencies in the family and then summing over loci. Let *F* be the family from parents *Pk* × *P*_*m*_ (parent indices *k, m*). Let *j* index a diploid, biallelic locus with alleles *A* and *a*, and let the total number of loci be *L*. Let *a*_*j*_ and *d*_*j*_ denote the additive and dominance effects at locus *j*, estimated under the marker codings *M*[*i, j*] ∈ {0, 1, 2} (count of allele *a*: *AA*=0, *Aa*=1, *aa*=2) and *W*[*i, j*] ∈ {0, 1} (heterozygote indicator: *W*=1 for *Aa*, else 0). Let *PAA*_*F,j*_, *PAa*_*F,j*_, and *Paa*_*F,j*_ be the genotype frequencies in *F* at locus *j* (with *PAA*_*F,j*_ + *PAa*_*F,j*_ + *Paa*_*F,j*_ = 1). The expected single-locus contribution is:

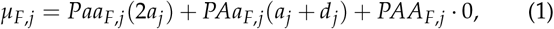

Summing across all *L* loci gives the family mean:

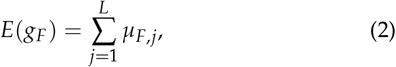

Progeny genotypic frequencies after any number of selfing cycles can be obtained with the following general matrix notation:

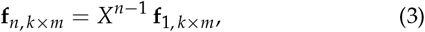

where **f**_1, *k*_*×*_*m*_ is a vector of F1 progeny conditional probabilities for the cross *P*_*k*_ ×*P*_*m*_, *X* is a column-stochastic square selfing transition matrix with progeny conditional probabilities in the rows and parental-genotype classes in the columns, and **f**_*n, k*_ × _*m*_ is the corresponding vector after *n* selfing cycles (see Appendix 1: Building the selfing transition matrix for diploids for more details).

### Prediction of within-family variance

For a single bi-allelic locus:

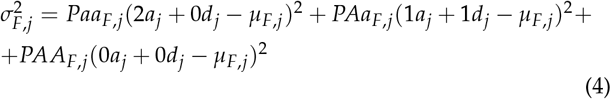

Where 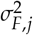 is the within-family variance for locus *j* and *µ*_*F,j*_ is calculated from Equation 1.

Considering the covariances between loci and dependence of additive and dominance marker effects, the within-family variance across all loci for a family is computed by:

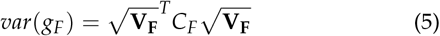

where 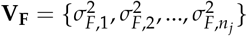 are locus-specific variances from Equation 4 and *C*_*F*_ is the correlation matrix of genotypic effects between loci (Fernandez-Gonzalez *et al*. 2025).

### Proportion of additive standard deviation lost

Controlling genetic diversity requires metrics that are both interpretable and robust to incomplete information, such as pedigree records. Genomic relationships are attractive because they capture relatedness due to recent and distant common ancestors (Keller et al. 2011). We developed the *proportion of additive standard deviation lost* (PropSD), which is dependent on genomic relationship matrices. PropSD is intuitive to tune, directly proportional to expected genetic gain, and applicable across breeding schemes (Fernandez-Gonzalez et al. 2025).

Let 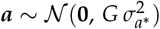 represent additive genetic values for the full parental pool of size *n*, and 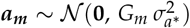 the additive values for the *n*_*m*_ selected parents. Selection reduces genetic dispersion, so the standard deviation among selected parents, 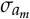, is typically smaller than that of the full pool, *σ*_*a*_. Hence, PropSD is:

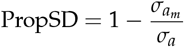

Although *σ*_*a*_ and 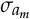 are unknown, we estimate them as the expected sample standard deviations from the corresponding multivariate distributions: 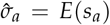 and 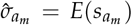. We first obtain the expected sample variances (with the unbiased denominators *n* − 1 and *n*_*m*_ − 1):

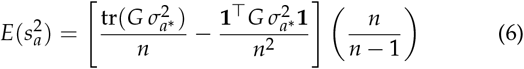

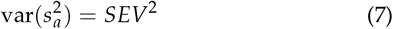

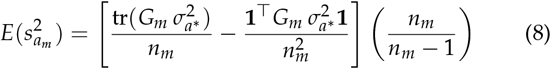

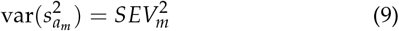

where *SEV*^2^ is the standard error of the variance computed as in Fernández-González and Isidro y Sánchez (2025) from the distribution 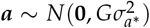 and 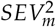 is the standard error of the variance corresponding to 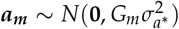.

Given that the sample variance follows a gamma distribution:

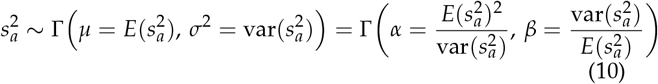

where *α* and *β* are the shape and scale parameters of a gamma distribution.

Using the probability density function of a gamma distribution, we can compute *E*(*s*_*a*_) as the moment of order 1/2. The pdf is:

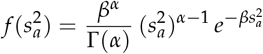

where Γ(·) denotes the gamma function. Gamma distributions satisfy Γ(*α, β*) = *β* · Γ(*α*, 1). Taking *β* outside lets us write:

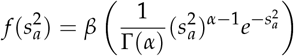

Define 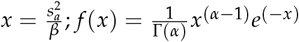. Applying the formula for the moment of order 1/2 of a continuous distribution:

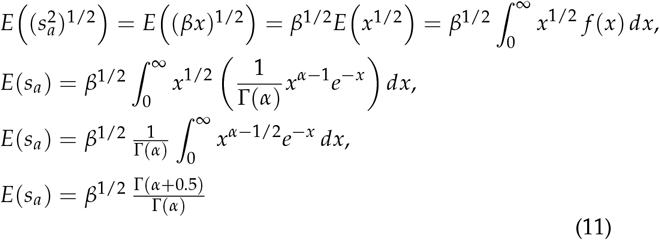

Applying the same steps to 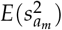 and 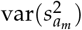 yields 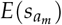. Therefore, the estimated proportion of standard deviation lost is:

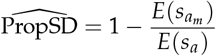

### Importance of the usefulness criterion

The usefulness criterion (UC), first conceptualized by Schnell and Utz (1975), is the linear combination of family mean and within-family variance:

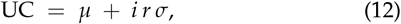

where *µ* is the family mean, *i* is the within-family selection intensity, *r* is the within-family selection accuracy, and *σ* is the within-family genetic standard deviation.

The contribution of the variance term depends on how much family standard deviations vary across families relative to the variation in family means, i.e., on the ratio var(*σ*)/ var(*µ*). A convenient summary of UC’s added value is the correlation *ρ* = cor(*µ*, UC): if *ρ* ≈ 1, UC is effectively equivalent to selection on *µ* alone; if *ρ* is relatively small, incorporating *σ* through UC becomes more beneficial.

To derive *ρ*, assume *µ* and *σ* are independent across families, so that cov(*µ, irσ*) = 0. Then:

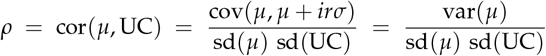

Using var(*a* + *b*) = var(*a*) + var(*b*) for independent *a, b* and var(*cX*) = *c*^2^ var(*X*), where c is a scalar and X any random variable,

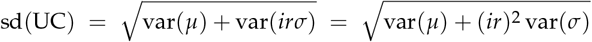

Therefore,

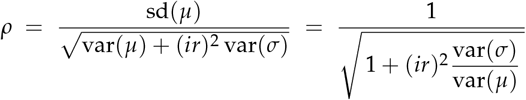

In summary, the correlation between *µ* and UC is:

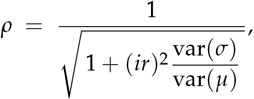

which depends on *i, r*, and the across-family variance ratio var(*σ*)/ var(*µ*).

To express the practical impact on expected response, define the **Usefulness Importance (UI)** as the multiplicative gain in response when moving from selection on *µ* to selection on UC, i.e., (genetic gain using UC) = UI × (genetic gain using *µ*). If the true target is UC, selecting on *µ* reduces accuracy by a factor *ρ*, and, by the Breeder’s equation, reduces expected genetic gain by the same factor. Hence:

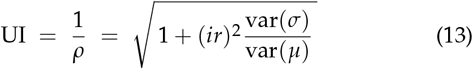

As an illustrative example, consider protein content in wheat, for which a ratio of var(*µ*)/var(*σ*) = 2.5 has been reported by Lado *et al*. (2017). Under a within-family selection intensity of *i* = 2 and a selection accuracy of *r* = 0.5, the correlation between the usefulness criterion (UC) and the family mean is *ρ*_*µ*_,*UC* ≈ 0.85. The corresponding usefulness importance is *𝒰I* = 1/*ρ*_*u*_*μ*C,𝒰C ≈ 1.176, indicating that selection based on UC would be expected to increase the response to selection by approximately 17.6% per cycle relative to selection on family means alone. Using Equation 13, Figure 2 illustrates the general relationship between var(*µ*)/var(*σ*) and both *ρ*_*µ*_,*UC* and its inverse (UI), across a range of within-family selection accuracies.

**Figure 2.**
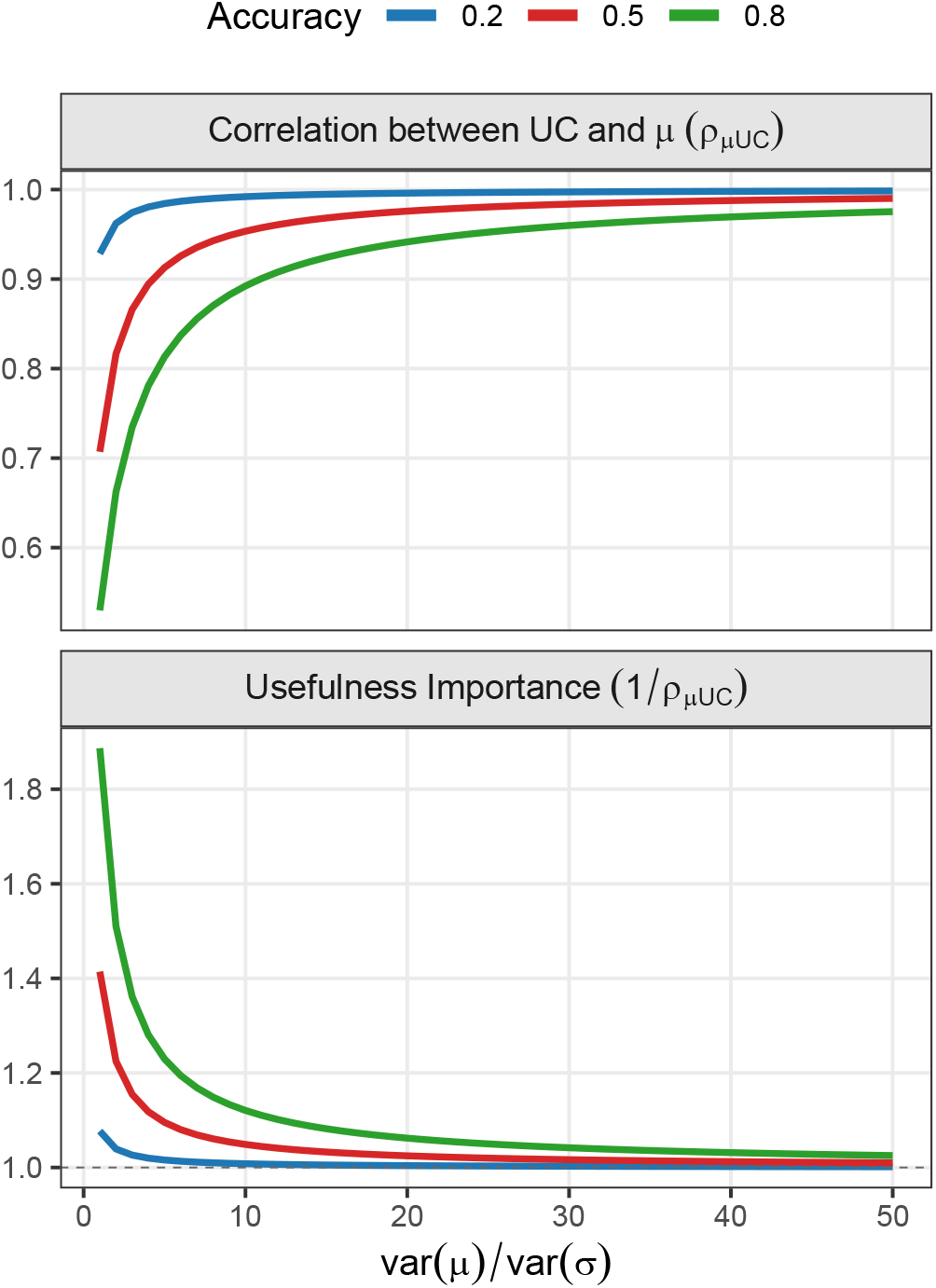
Relationship between var(*µ*)/var(*σ*) (variance of family means relative to the variance of family genetic standard deviations) and (top) the correlation between the usefulness criterion (UC) and the mean (*ρ*_*µ*_,*UC*), and (bottom) its reciprocal, *usefulness importance* (1/*ρµ,UC*). Curves correspond to three levels of within-family selection accuracy (0.2, 0.5, and 0.8), with a within-family selection intensity fixed at 2.

## Methods

We conducted stochastic simulations of two breeding programs (line and hybrid). For each breeding program, we simulated two control scenarios: conventional phenotypic selection (PS) and GS, with random mating of parents. In addition to eight scenarios with genomic mating based on different criteria, with and without genetic diversity control. An overview of the simulated scenarios and the factors they consider is shown in Table 1. For OGM, we used the `SimpleMating` (Peixoto *et al*. 2024a) and `MateR` (Fernandez-Gonzalez *et al*. 2025) packages, which both implement methods to predict within-family variance under different assumptions, with or without dominance, as required in this study.

**Table 1.** Description of simulated scenarios and the factors considered by each (family mean, within-, and between-family variance)

| Scenario | Description | Family mean | Within-family variance | Between-family variance<sup>a</sup> |
| --- | --- | --- | --- | --- |
| PS_RM | Phenotypic selection with random mating | – | – | – |
| GS_RM | Genomic selection with random mating | – | – | – |
| GS_FM | Genomic selection with mating based on family mean | ✓ | – | – |
| GS_UC | Genomic selection with mating based on usefulness criterion | ✓ | ✓ | – |
| 2F-SM | Two factor optimal genomic mating with SimpleMating | ✓ | – | ✓ |
| 2F-MtR | Two factor optimal genomic mating with MateR | ✓ | – | ✓ |
| 2F-MtR+OR | Two factor optimal recycling and genomic mating with MateR | ✓ | – | ✓ |
| 3F-SM | Three factor optimal genomic mating with SimpleMating | ✓ | ✓ | ✓ |
| 3F-MtR | Three factor optimal genomic mating with MateR | ✓ | ✓ | ✓ |
| 3F-MtR+OR | Three factor optimal recycling and genomic mating with MateR | ✓ | ✓ | ✓ |

<sup>a</sup> A checkmark (✓) indicates the factor is included; “–” indicates it is not.

### Description of the breeding programs

#### Line

The line breeding program followed established wheat breeding schemes producing doubled haploid (DH) lines (Gaynor *et al*. 2017; Bančič *et al*. 2025). Each cycle began with inbred parental line crosses to create an F_1_ population, from which DH lines were derived and evaluated in multi-environment, multi-year trials. Each cycle released a pure-line variety exploiting additive variance.

#### Hybrid

The hybrid breeding program simulated maize DH line development within heterotic pools (Bernardo 2009; Powell *et al*. 2020; Bančič *et al*. 2025). Parental line crosses produced F_1_ populations from which DH lines were generated and testcross-evaluated for general (GCA) and specific combining ability (SCA). Each cycle resulted in an F_1_ hybrid exploiting additive and dominance variance, propagated through repeated crosses of its inbred parents.

### Genome simulation

Genome simulations were performed using `AlphaSimR` with backward-in-time coalescent models (Chen *et al*. 2009; Gaynor *et al*. 2021). Details are presented below for each breeding program:

#### Line

We simulated a 14-chromosome genome representing tetraploid durum wheat (*Triticum durum*), with genetic and physical lengths of 1.43 Morgans and 8 × 10_8_ base pairs. Recombination and mutation rates were 1.8 × 10−9 and 2 × 10−9 per base pair. Effective population size (*Ne*) followed wheat domestication dynamics (Thuillet *et al*. 2005; Peng *et al*. 2011), increasing from an initial *N*_*e*_ of 50 to 1,000 at 100 generations ago, 6,000 at 1,000 generations ago, 12,000 at 10,000 generations ago, and reaching 32,000 at 100,000 generations ago.

#### Hybrid

We modeled maize (*Zea mays*) using 10 chromosomes (2 Morgans, 2 × 108 bp each). Recombination and mutation rates were 1.25 × 10−8 and 1 × 10−8 per bp, with *N*_*e*_ = 100 per heterotic group, consistent with maize history (Hickey *et al*. 2014; Powell *et al*. 2020). The two heterotic groups were split 30 generations ago to reflect the historical public–private divergence reported for sweet corn genotypes in the United States (Peixoto *et al*. 2024b).

### Simulation of founder genotypes

Founders were created by sampling chromosome pairs from simulated genomes. For each breeding program, biallelic QTNs and SNPs were drawn uniformly at random per chromosome at the counts specified below.

#### Line

We assembled 50 diploid founders in Hardy–Weinberg equilibrium, sampling 14 chromosome pairs per genotype. For the trait architecture, we selected 1,000 QTNs and 400 SNPs per chromosome (14,000 QTNs; 5,600 SNPs total).

#### Hybrid

For each heterotic group, we assembled 50 diploid founders by sampling 10 chromosome pairs per genotype. Each chromosome carried 300 QTNs and 1,250 SNPs (3,000 QTNs; 12,500 SNPs total).

### Simulation of genetic values

Genetic values for yield were computed as the sum of locus effects over the QTNs specified for each breeding program. For every effect class included, additive (*a*), dominance (*d*; when relevant), and genotype-by-year interaction (*G* × *Y*), raw effects were sampled from a standard normal and rescaled to the program-specific target variances indicated below. `AlphaSimR` trait types are given in parentheses.

#### Line

(AG in `AlphaSimR`) At 14,000 QTNs, additive effects were scaled to 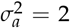 in the founder population, and *G* × *Y* effects were scaled to match (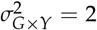).

#### Hybrids

(ADG in `AlphaSimR`) At 3,000 QTNs, additive effects were scaled to 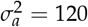, with *G* × *Y* effects matched accordingly 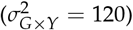.

Dominance effects (*d*) were derived as the product of the absolute additive effect (|*a*_*i*_|) and a locus-specific dominance degree (*δ*_*i*_). The dominance effect of QTN *i* was calculated as:

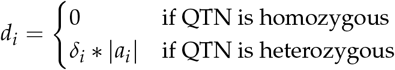

Dominance degrees were sampled from a normal distribution with mean *µδ* = 0.92 and variance 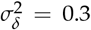, consistent with historical heterosis in maize (Troyer and Wellin 2009; Powell et al. 2020; Peixoto et al. 2024b).

### Simulation of phenotypes

We simulated phenotypes by adding random error terms to the genetic values. These error terms followed a normal distribution with mean zero and variance adjusted to target the specified heritability (*h*_2_) at each testing stage. Entry mean narrow-sense heritability increased from early (unreplicated) to late (replicated) testing stages, calculated as:

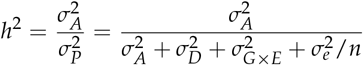

where *n* is the number of replications per genotype, 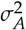 is additive genetic variance, 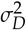 is dominance variance, 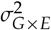 is genotype-by-environment interaction variance, 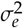 is residual variance, and 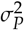 is total phenotypic variance.

### Recent (burn-in) breeding phase

#### Line

We simulated a conventional PS program for 20 years (Figure S1). Each year started with 100 biparental crosses randomly chosen without replacement from 1,225 possible parental combinations, generating 10,000 doubled haploid (DH) lines (100 per family).

In year three, we planted the DH lines in a headrow nursery (HRW; *h*_2_ = 0.18), selecting the top five lines per family (500 total). In year four, selected lines underwent preliminary yield trials (PYT; *h*_2_ = 0.33), advancing the top 50 lines. In year five, advanced yield trials (AYT; *h*_2_ = 0.67) identified the best ten lines. Finally, in year six, the ten lines were evaluated in elite yield trials (EYT; *h*_2_ = 0.8) and recycled to replace the ten oldest parents.

#### Hybrid

We simulated a conventional PS program for 20 years (Figure S2). Each heterotic group annually underwent 80 biparental crosses randomly chosen from 1,225 combinations, generating 4,800 DH lines (60 per family).

DH lines were crossed with a single tester from the opposite heterotic group, evaluating F_1_ hybrids in test-cross trial 1 (TC1; *h*_2_ = 0.3). We selected the top four inbreds per family (320 total), crossing each with three testers in year three for test-cross trial 2 (TC2; *h*_2_ = 0.47). From these, we chose 40 inbreds based on general combining ability (GCA). In year four, 40 selected inbreds crossed with five testers were evaluated in test-cross trial 3 (TC3; *h*_2_ = 0.64), selecting 20 hybrids based on specific combining ability (SCA).

In year five, hybrid yield trial 1 (HYT1; *h*_2_ = 0.78) evaluated 20 hybrids, selecting four. The inbred parents of the top ten hybrids replaced the oldest parents. In year six, hybrid yield trial 2 (HYT2; *h*_2_ = 0.9) identified the best hybrid, whose inbred parent replaced the oldest tester and advanced to commercial release.

### Future breeding phase (control scenarios)

We simulated two control scenarios (PS and GS) over 30 years for each breeding program, performing random parental crosses in each. The control PS scenarios retained the structure of the previously described burn-in phases. Here, we describe only the GS scenarios.

#### Line

Each year began with 100 biparental crosses randomly selected without replacement from the 1,225 possible combinations of 50 parental lines, generating 10,000 DH lines (100 per family) (Figure 3). DH lines were genotyped, and their genomic estimated additive values (GEAVs) were predicted from a GS model trained on the preceding five years of yield trial data (PYT, AYT, EYT). Training data were updated annually by replacing the oldest records with new results, maintaining a fixed training set size.

**Figure 3.**
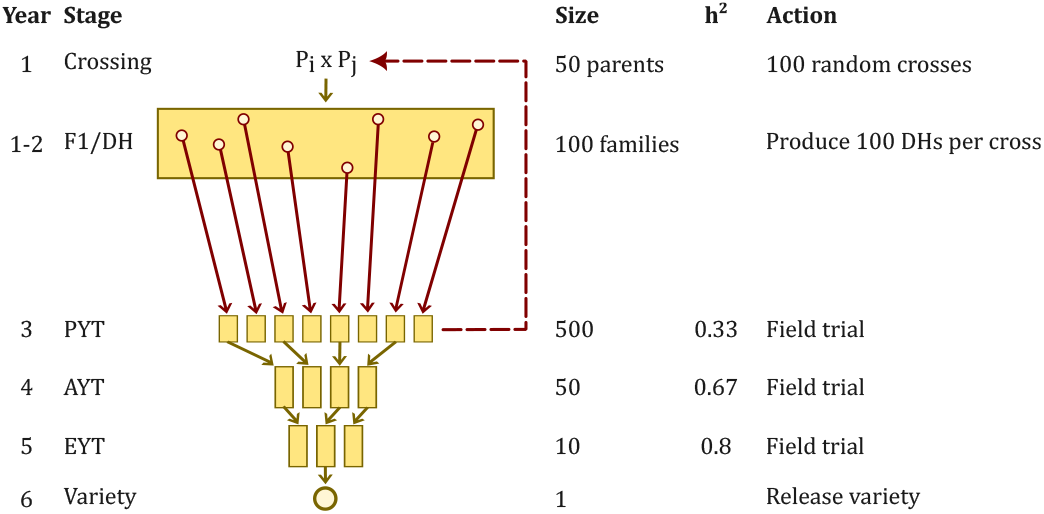
Schematic overview of the line-breeding genomic selection (GS) program using double haploid (DH) technology (adapted with modifications from Gaynor *et al*. (2017); Bančič *et al*. (2025)). The program consists of key breeding stages: `DH`, `PYT` (preliminary yield trial), `AYT` (advanced yield trial), and `EYT` (elite yield trial). Each stage includes varying numbers of individuals (`Size`), yield heritability (*h*2), and specific actions. The crimson red dashed line indicates the recycling of individuals from `PYT` into the next breeding cycle. All actions shown in crimson red highlight changes introduced by GS, as described in Future breeding phase (control scenarios).

From the 10,000 DH lines, we selected the top five per family (500 total) based on GEAVs. These were evaluated in preliminary yield trials (PYT; *h*_2_ = 0.33), advancing the top 50 phenotypically performing lines. Concurrently, the top 10 lines (based on GEAVs) replaced the oldest parents in the crossing block. Subsequent phenotypic selection stages (AYT and EYT) proceeded identically to the burn-in phase.

##### Hybrid

Annually, we randomly selected 80 biparental crosses per heterotic group from 1,225 possible combinations, producing 4,800 DH lines (60 per family) (Figure 4). We genotyped DH lines and predicted their hybrid progeny performance (general combining ability, GCA; specific combining ability, SCA) using a GS model trained on two recent years of test-cross (TC2, TC3) and hybrid yield (HYT1) trial data. Training data were updated annually.

**Figure 4.**
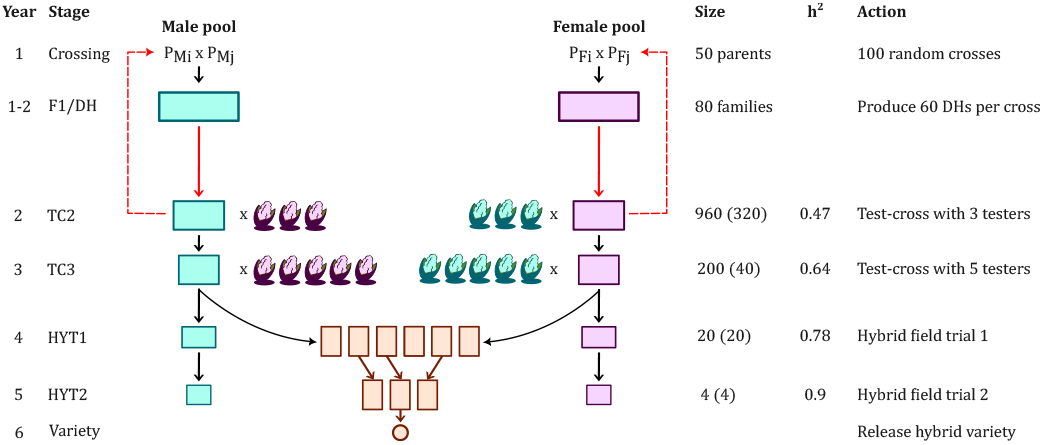
Schematic overview of the hybrid-breeding genomic selection (GS) program using double haploid (DH) technology (adapted with modifications from Bernardo (2009); Powell *et al*. (2020); Bančič *et al*. (2025)). The program consists of key breeding stages: `DH`, `TC` (test-cross trial), and `HYT` (hybrid yield trial). Each stage includes varying numbers of individuals (`Size`), yield heritability (*h*^2^), and specific actions. The population size at each `TC` and `HYT` stage refers to the number of hybrids in each heterotic pool, while values in parentheses indicate the number of DH inbreds in each pool. The red dashed line represents the recycling of individuals from `TC2` into the next breeding cycle. All actions shown in red highlight changes introduced by GS, as described in Future breeding phase (control scenarios).

We selected the top four DH inbreds per family (320 total) based on predicted hybrid performance. These inbreds were crossed with three testers from the opposite heterotic group, and the resulting hybrids were evaluated in TC2 trials (*h*_2_ = 0.47). The 40 inbreds with the highest GCA advanced to later stages, while the top 10 inbreds, ranked by their predicted hybrid genotypic values with five testers, replaced the oldest parents in the crossing block. All subsequent selection stages (TC3, HYT1, HYT2) followed the same structure as described in the burn-in phase.

### Future breeding phase (genomic mating scenarios)

Genomic mating was implemented by modifying the GS control schemes at the crossing block: parents were mated non-randomly according to the criterion specific to each scenario. As described in Table 1, we evaluated eight scenarios (`GS_FM`, `GS_UC`, `2F-SM`, `3F-SM`, `2F-MtR`, `3F-MtR`, `2F-MtR+OR`, and `3F-MtR+OR`).

In `GS_FM`: crosses were prioritized by their predicted family mean (Prediction of family mean); for hybrid breeding, the family mean was the mean predicted progeny performance when crossed to five testers from the opposite heterotic pool. `GS_UC`: crosses were selected based on their UC (Eq. 12); within-family variance was predicted as in Prediction of within-family variance.

In `2F-SM` and `3F-SM`: `SimpleMating` was used to optimize mating plans that balance expected gain with targeted inbreeding (relatedness among crosses); predictions of family mean and within-family variance followed Peixoto *et al*. (2024a).

In `2F-MtR` and `3F-MtR`: `MateR` was used to optimize mating, balancing expected gain with target genetic diversity (PropSD) thresholds as described in Proportion of additive standard deviation lost; family means and within-family variances followed Prediction of family mean and Prediction of within-family variance. `2F-MtR+OR` and `3F-MtR+OR`: identical to the latter `MtR` scenarios but recycle the *optimal* individuals that balance gain and variance, rather than the top performers (Figure 1).

### Tuning genetic diversity parameters

For `SimpleMating`, diversity was controlled through the covariance between parental pairs, obtained from *G* relationship matrices (Kinghorn 2011; Peixoto et al. 2024a). We tuned it by retaining different proportions of related crosses, ranging from 10% to 90%, with 10% representing the strictest diversity constraint. The minimum individual contribution was set to 0 and the initial maximum to 10. If no feasible solution was identified under this constraint, the maximum contribution limit was incrementally increased until a valid solution was found.

For `MateR` (Fernandez-Gonzalez et al. 2025), genetic diversity was controlled by tuning the PropSD metric (see Proportion of additive standard deviation lost section) for values ranging from 0.5% to 10% additive standard deviation lost, with 0.5% being the strictest diversity constraint. PropSD thresholds set to control diversity in the recycling stage were equal to those used to optimize mating in the crossing block. No constraints were imposed on the minimum and maximum number of participations per individual.

### Genomic selection modeling

We applied genomic best linear unbiased prediction (gBLUP) models tailored to breeding programs, specific genetic effects (additive and/or dominance), and analytical objectives (marker effects or within-family prediction accuracy). Models were fitted using `sommer` v4.3.7 (Covarrubias-Pazaran 2016).

#### Additive model (A)

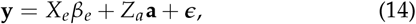

where **y** is a vector of phenotypic entry means; *X*_*e*_ is the incidence matrix linking observations to fixed year effects (*β*_*e*_); *Z*_*a*_ links observations to additive genetic effects (**a**), with 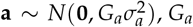, is the additive genomic relationship matrix and 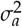 is the additive genetic variance; and 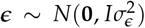 is the vector of residual errors with variance 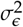.

#### Additive and dominance model (A+D)

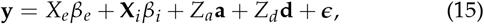

where, in addition to terms described previously, **X**_*i*_ is the vector of inbreeding values, (*β*_*i*_) is a fixed slope for inbreeding, capturing variation due to different inbreeding levels (Endelman 2023); and *Z*_*d*_ links observations to dominance genetic effects 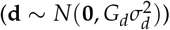, where *G*_*d*_ is the dominance genomic relationship matrix and 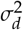 is the dominance genetic variance.

### Comparative analysis

We compared our different simulated scenarios in terms of realized genetic means, genetic variance, genic variance, inbreeding rate, and the efficiency of converting genetic diversity into gain.

The comparison was performed by fitting a mixed model where scenarios are set as fixed and replicates as random effects.

The true genetic means, in addition to the total genetic and genic variability, were obtained using `AlphaSimR` built-in functions (`meanG`, `varG`, and `genicVarG`). Average group coancestry and inbreeding rate were estimated from pedigree and genomic information in the parental pool(s). The *A* matrix was obtained using the `AGHmatrix` package (Amadeu *et al*. 2023), and *G* was computed following (VanRaden 2008):

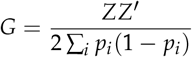

where *Z* is the centered genotype matrix where each element *Z*_*ij*_ = genotype_*ij*_ − 2*p*_*i*_ with genotype_*ij*_ coded as 0, 1, or 2 for individual *j* at marker *i*, and *p*_*i*_ is the reference allele frequency at marker *i* in the base population.

The efficiency of converting genetic diversity into genetic gain was measured following Gorjanc *et al*. (2018), by regressing the standardized genetic gain achieved,

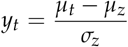

on lost genic standard deviation,

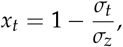

i.e.

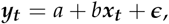

where *µ* is the genetic mean, *σ* is the genic standard deviation, *b* is the conversion efficiency, *z* is the initial year of the future breeding phase, and a data-point is available each later year *t. a* and *b* are a fixed intercept and an slope respectively, and ***ϵ*** is a vector of i.i.d. residuals.

To ensure a fair comparison between `SimpleMating` and `MateR`, we investigated the Pareto front of the realized genetic gain-variance trade-offs across all simulated programs. An optimal solution was selected as representative for subsequent analyses based on the best trade-off in the last year (30).

### Software implementation

All simulations were conducted using `AlphaSimR` v1.6.0 (Gaynor *et al*. 2021). Subsequent data analyses and visualizations were performed in R v4.3.2 (R Core Team 2023), primarily with the `ggplot2` package (Wickham 2016). Optimal genomic mating was implemented with `SimpleMating` v0.1.9001 (Peixoto *et al*. 2024a) and `MateR` v1.0.0 (Fernandez-Gonzalez *et al*. 2025). Optimal recycling was also implemented through `MateR` v1.0.0, powered by the `TrainSel` v3.0 algorithm (Akdemir *et al*. 2021).

## Results

### Gain-variance Pareto front

Figure 5 illustrates the range of realized gain–variance trade-offs achieved under various diversity control thresholds for the two used packages (`SimpleMating` and `MateR`) and tested strategies implemented in `MateR` (OGM with and without optimal recycling). Overall, GS with random mating (`GS_RM`) was good at preserving diversity, but inferior for gain. In contrast, truncation selection of crosses based on family mean (`GS_FM`) or UC (`GS_UC`) reached relatively higher gains at the expense of depleting the most variance. Meanwhile, the OGM scenarios (`SM`, `MtR`, and `MtR+OR`) achieved better optimal solutions, with comparable or higher gains that deplete less variance.

**Figure 5.**
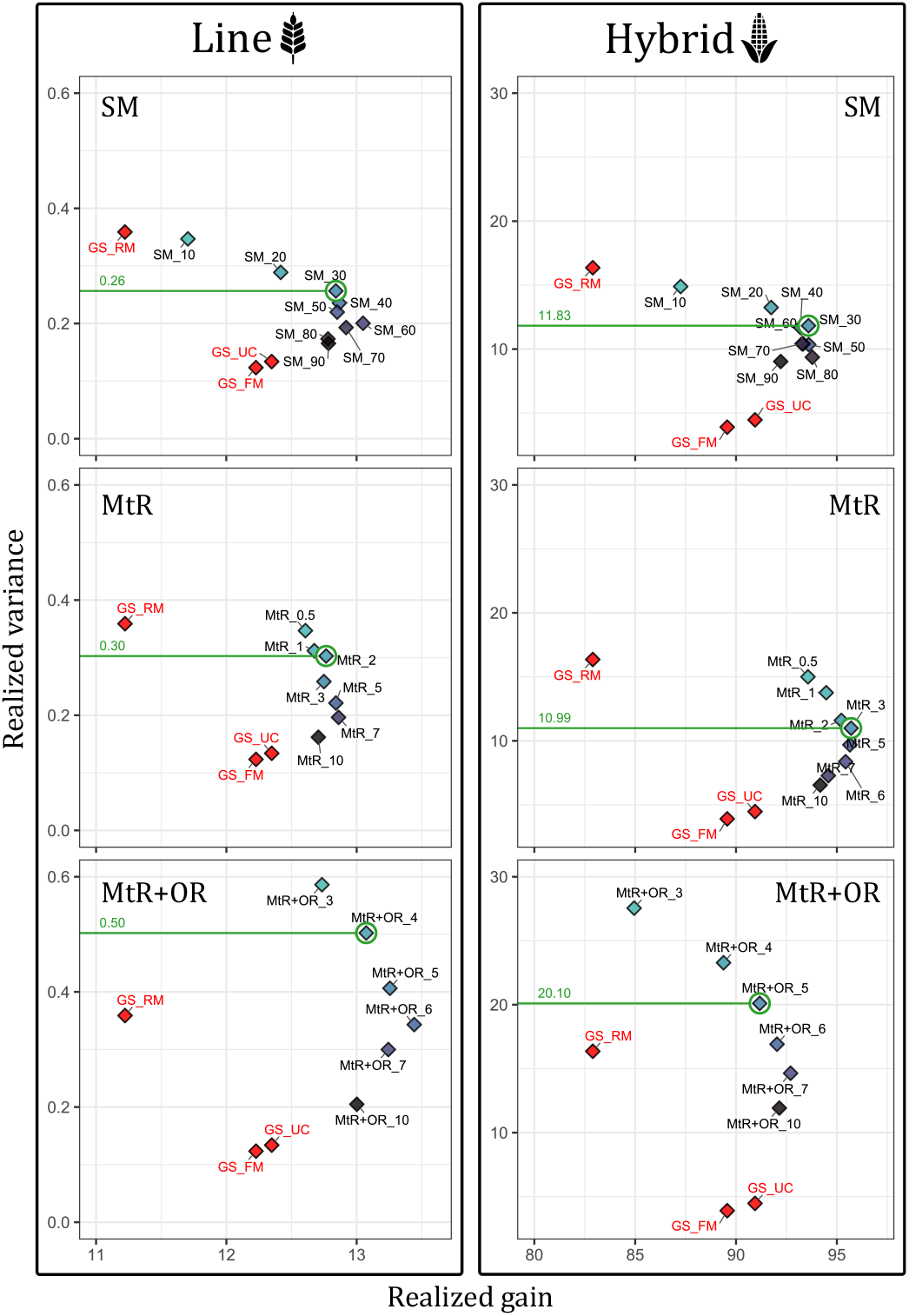
Realized genetic gain and variance in the last year (30) for line and hybrid breeding programs. Each point represents the mean of 100 simulation replicates. The points in red denote the following genomic selection scenarios: random mating (`GS_RM`), mating based on family mean (`GS_FM`), and usefulness criterion (`GS_UC`). The remaining points are the different genetic diversity control thresholds for different packages and breeding strategies, ranging from diversity-strict to more lenient thresholds: 10% to 90% for `SimpleMating` (`SM`), 0.5% to 10% for `MateR` (`MtR`), and 3% to 10% for `MateR+OR` (`MtR+OR`) (see Tuning genetic diversity parameters). The color gradient from cyan to dark purple represents the increasing leniency in diversity control. The green circle and line highlight the Pareto optimal scenario for each strategy and package, with their corresponding realized variance.

In line and hybrid programs, the Pareto optimal solution was to retain 30% of related crosses for `SimpleMating`, setting PropSD at 2–3% for OGM only with `MateR`, and 4–5% when optimal recycling is implemented with OGM through `MateR` (`MtR+OR`). Integrating optimal recycling and genomic mating resulted in the expansion of the Pareto with dominating solutions relative to genomic mating alone (in line breeding), in addition to random and truncation genomic mating scenarios in both programs. Interestingly, `GS_RM` was only dominated in terms of both, variance and gain with the integrated strategy. The optimal solutions of the integrated strategy at 4–5% PropSD allowed for saving about 1.6–2.0 times the variance retained by OGM alone (using `SimpleMating` `and` `MateR`) and 3.5–5.0 times the variance kept by truncation mating scenarios, without compromising long-term gains relative to `GS_FM` and `GS_UC` in both programs (Table S2). Separate trends of genetic mean, genetic variance, and genic standard deviation with optimal diversity control thresholds are shown in Figure S3.

### Conversion efficiency

Figure 6 and Table S1 summarize the conversion efficiency of genetic diversty into genetic gain for each scenario in both breeding programs. Generally, phenotypic selection with random mating (`PS_RM`) achieved the highest conversion efficiency. However, it delivered around 40% and 30% lower standardized gains in line and hybrid programs, relative to the least efficient truncation mating scenarios (`GS_FM` and `GS_UC`). Scenarios that implement OGM only, using `SimpleMating` and `MateR`, achieved efficiencies comparable or higher than those of genomic selection with random mating (`GS_RM`), and both were more efficient than truncation mating ones.

**Figure 6.**
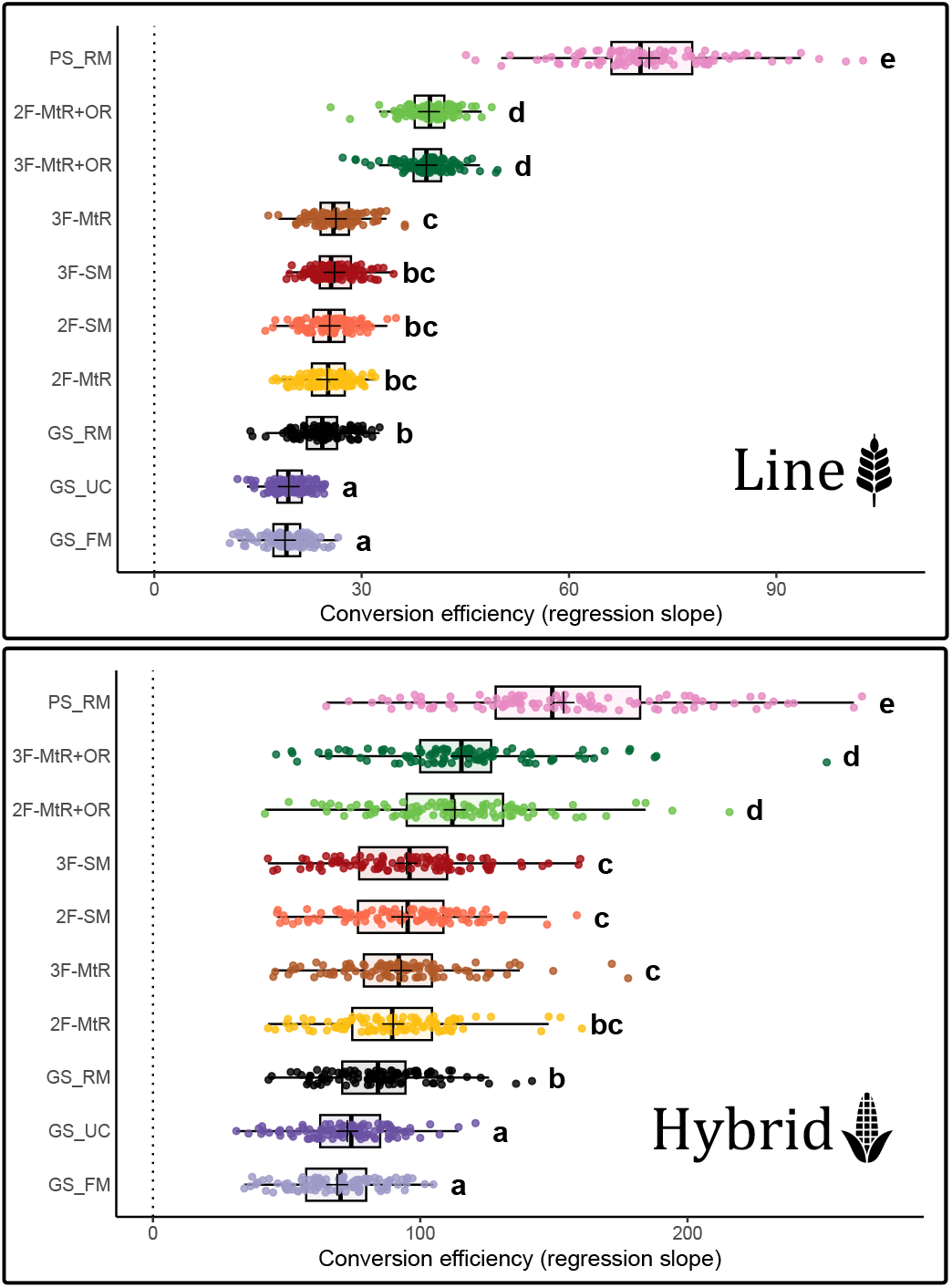
The conversion efficiency of genic standard deviation into genetic gain for each simulated scenario in the line and hybrid breeding programs. All scenarios are replicated 100 times and described in Table 1. Scenarios with the same letter do not differ significantly (Tukey’s HSD test, 99% confidence interval).

The integrated strategy of optimal recycling and genomic mating (`2F-MtR+OR` and `3F-MtR+OR`) consistently and significantly improved conversion efficiency relative to all simulated genomic strategies, reducing the gap between them and `PS_RM`. Both were around 49.8% more efficient in line breeding compared to the best genomic mating only scenario (21.6% in hybrid), and around 108.2% more efficient when compared to `GS_FM` (67.4% in hybrid). There were no significant differences between the scenarios that accounted for within-family variance (`3F`) and their counterparts (`2F`).

### Average group coancestry and inbreeding rate

We were interested to know how our different simulated strategies compare to the inbreeding rate (Δ*F*). Hence, we recorded the average group coancestry using pedigree and genomic information during the future breeding phase in the parental pool(s) to measure the realized Δ*F* (Figure 7 and Table S3).

**Figure 7.**
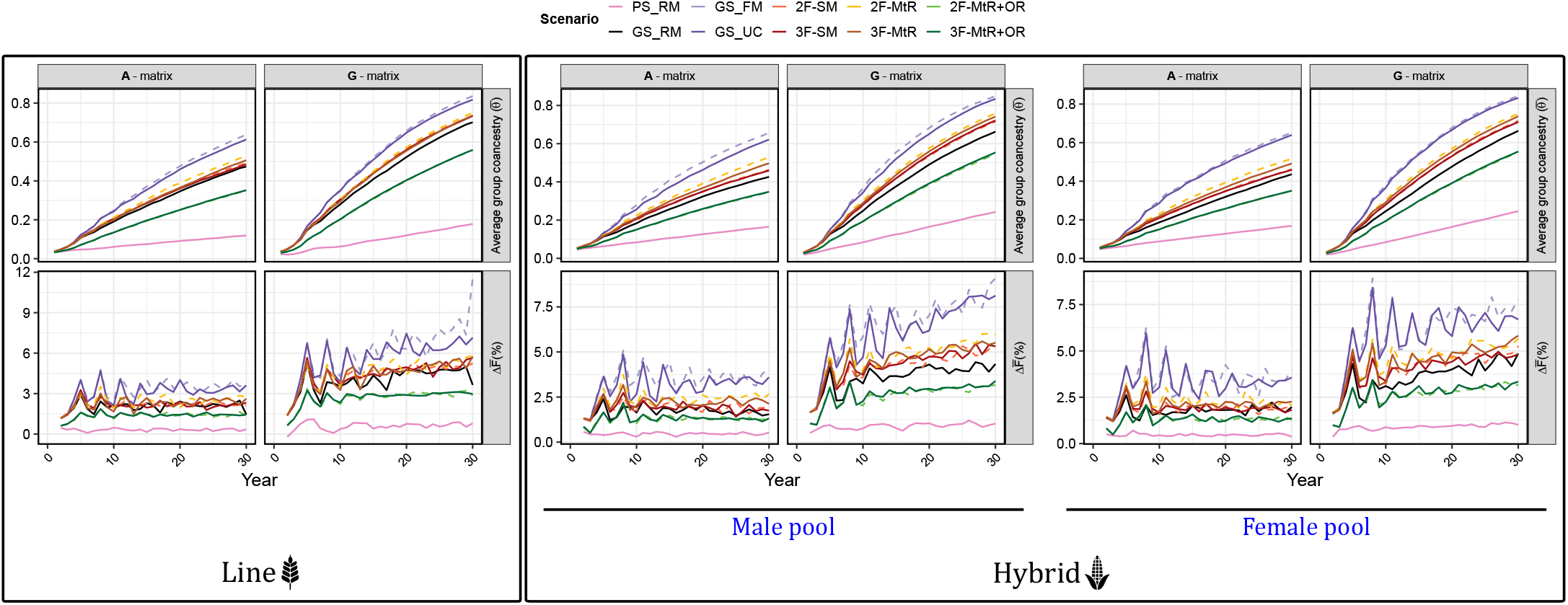
The average group coancestry and realized inbreeding rate recorded in the parental pools during the future breeding phase of the line and hybrid breeding programs. All scenarios are replicated 100 times and described in Table 1.

Inbreeding rates estimated from *A* and *G* matrices (Δ*F*_*A*_ and Δ*F*_*G*_) were generally consistent and positively correlated across scenarios and programs, with Δ*F*_*G*_ tending to yield slightly higher and more variable values as it captures realized Mendelian sampling and drift effects beyond pedigree expectations. Among all scenarios, phenotypic selection with random mating (`PS_RM`) achieved the lowest realized Δ*F* across both line and hybrid programs, with Δ*F*_*A*_ not exceeding 0.5% and Δ*F*_*G*_ not exceeding 1.1%. In contrast, the highest inbreeding rates were realized by the truncation mating strategies (`GS_FM` and `GS_UC`), which reached approximately 3–3.5% for Δ*F*_*A*_ and 6% for Δ*F*_*G*_.

Genomic selection with random mating (`GS_RM`) represented the second lowest inbreeding rate scenario, with Δ*F*_*A*_ ranging from about 1.7–2.0% and 3.6–4.0% for Δ*F*_*G*_ across programs. The optimal genomic mating scenarios without optimal recycling (`SM` and `MtR`) achieved slightly higher rates, with Δ*F*_*A*_ around 1.9–2.4% and Δ*F*_*G*_ around 4.1–4.7%. The implementation of optimal recycling together with genomic mating markedly reduced realized inbreeding rates relative to OGM alone. Across both programs, Δ*F*_*A*_ decreased to about 1.3–1.4% and Δ*F*_*G*_ to about 2.6–2.7%, corresponding to rates that were 1.5–1.8 times lower than those observed with genomic mating only and 2.2–2.6 times lower than those of truncation mating scenarios.

### Dynamics of variance ratio (family *µ* and *σ*_*a*_)

Figure 8 and Table S4 show the dynamics of the variance ratio between family genetic means (*var*(*µ*)) and family genetic standard deviations (*var*(*σ*_*a*_)) in line and hybrid programs. Initially, *var*(*µ*) greatly exceeded *var*(*σ*_*a*_), resulting in very high ratios of approximately 299 and 334 in the line and hybrid programs, respectively. Over successive breeding cycles, the ratio kept declining, reaching values near 60 in the line program and 135 in the hybrid program by year 30, as the variance in *µ* converged toward the variance in *σ*_*a*_. The mean difference in annual gains between the truncation mating scenarios (`GS_FM` and `GS_UC`) was close to zero with positive values, indicating a slight advantage in accounting for within-family variance.

**Figure 8.**
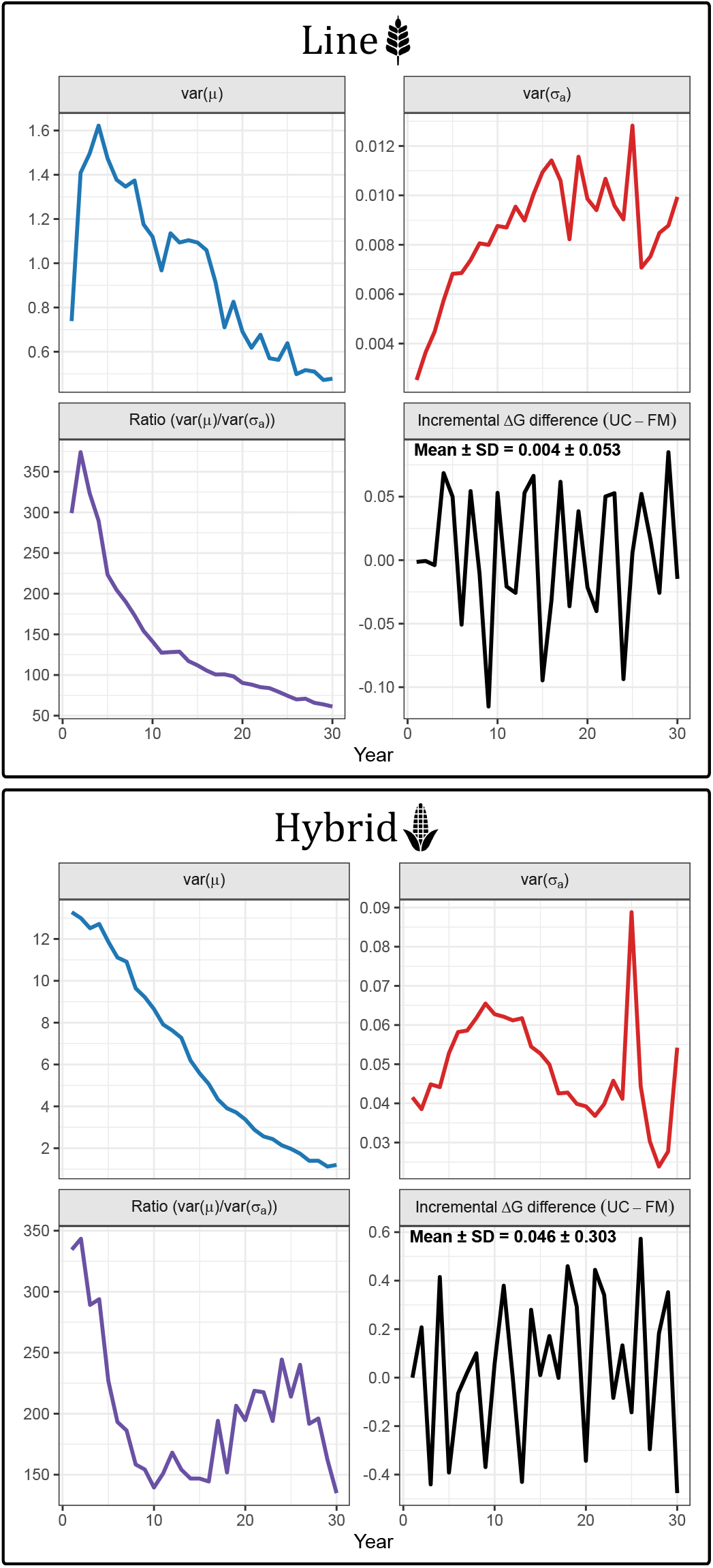
Dynamics of variance in family genetic means (*var*(*µ*)), family genetic standard deviations (*var*(*σ*_*a*_)), and their ratio 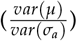 over 30 years of future breeding in line and hybrid programs. Lines represent means across 100 replicates. The lower-right panel shows the incremental Δ*G* difference, defined as the difference in annual genetic gain between the two truncation genomic mating scenarios (`GS_FM` and `GS_UC`). For the hybrid program, variances and ratios are averaged across male and female pools.

### Usefulness Importance

Attempting to explain the limited advantage of accounting for within-family variability, we monitored UI across future breeding years in both line and hybrid breeding programs. In general, UI increased as the ratio var(*µ*)/var(*σ*_*a*_) declined (Figures 9 and 8). However, average UI values ranged only from 1.003 to 1.005 in both programs, indicating that the advantage of UC relative to selection on family means alone was minimal and corresponded to an expected average additional gains of only 0.3–0.5% per cycle.

**Figure 9.**
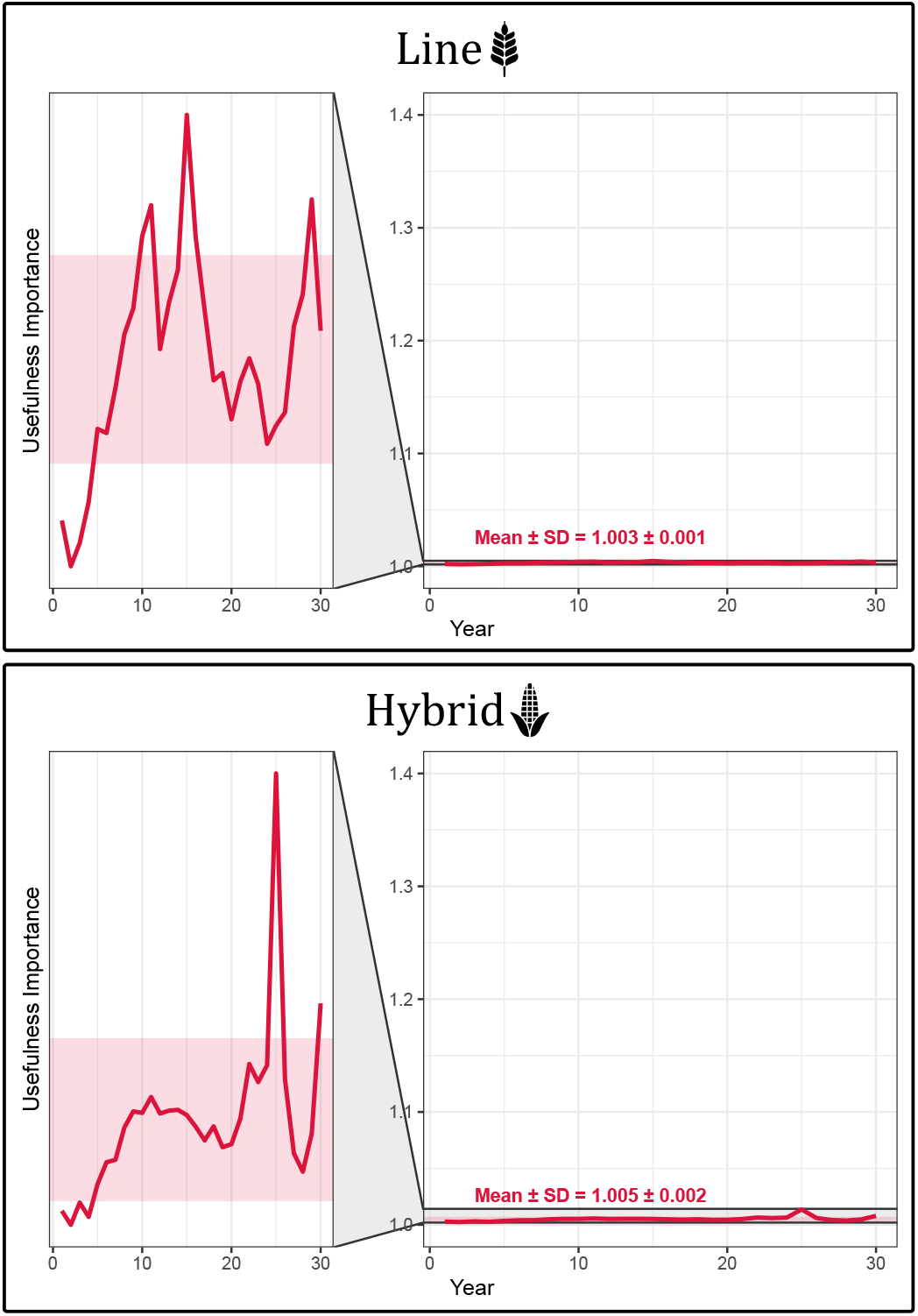
Trend of Usefulness Importance (UI) for the line and hybrid programs over the 30 years of future breeding. For the hybrid program, the trend is the average across male and female pools.

## Discussion

In this study, we evaluated an integrated strategy combining optimal recycling and genomic mating across two distinct breeding systems (line and hybrid), encompassing diverse trait architectures and genetic effects. We also presented the use of our newly developed metric (PropSD) to control genetic diversity in different breeding systems and stages (crossing block and recycling). Furthermore, we systematically dissected the factors that can be accounted for in the objective function of OGM to expose the relative importance of within-family variance.

### A step toward sustainable genomic selection programs

Crop breeding has long relied on phenotypic information to perform crosses and selection. Despite its practical limitations in terms of cost, cycle length, and suboptimal gains, it is very efficient in translating variance into gain due to the small amount of variability lost. Exploiting genomic information significantly enhances gains relative to PS programs; however, at the expense of a severe compromise to long term gains of relevant traits, because of the higher precision in selecting genomically related individuals (Crossa *et al*. 2017; Alemu *et al*. 2024). For this reason, there is a need to improve the efficiency of GS breeding programs.

Parental crosses are traditionally selected based on their predicted family mean performance or UC. That is a truncation mating approach in which crosses are sorted based on these criteria and the top *elite* × *elite* ones are picked, ignoring genetic diversity. This further exacerbates the negative impact of GS on the diversity of breeding populations. Hence, OGM was established to solve this problem, through finding a mating plan that balances the expected gain with population diversity, alleviating the adverse impact of truncation mating strategies in GS programs.

Here, we show that OGM alone is not sufficient to achieve our sustainable goal, a considerable amount of diversity is *leaked* in a different critical step of breeding schemes; the recycling step. Recycling can also be optimized; instead of selecting the top-performing individuals, we should find those who balance expected gains with variance (Figure 1). This aspect was not considered by previous studies aimed at genetic diversity preservation (Kinghorn 2011; Akdemir and Sánchez 2016; Gorjanc *et al*. 2018; Gorjanc and Hickey 2018; Danguy des Déserts *et al*. 2023; Sakurai *et al*. 2024; Peixoto *et al*. 2024b,a; Endelman 2025). Our results indicated that implementing an integrated strategy of optimal recycling and genomic mating can save double the amount of variability realized by OGM alone, and quadruple that of truncation mating strategies (Figure 5 and Table S2). This was equivalent to maintaining around 1.7 and 2.4 times less realized inbreeding rates (Figure 7 and Table S3).

### Improved conversion efficiency

We evaluated the efficiency of translating variance into gain and found that while OGM enhances it relative to truncation mating strategies, a wide gap remains between them and the most efficient scenario of PS (Figure 6). This was in agreement with the results of Gorjanc *et al*. (2018) and explains the success of their two-part breeding strategy with OGM in reaching an efficiency comparable to that of PS (Gaynor et al. 2017; Gorjanc et al. 2018). That improvement was driven by the increased gain due to the rapid recycling in the population improvement part while controlling diversity in the crossing block.

Combining optimal recycling with genomic mating in our work also improved conversion efficiency by around 21.6–49.8% compared to the sole implementation of OGM and 67.4–108.2% compared to truncation mating scenarios, approaching that of PS (Figure 6 and Table S1). We hypothesize that integrating optimal recycling and genomic mating in a two-part GS program could further expand the gain–variance Pareto front, due to the better control over genetic diversity. It could also be beneficial for the implementation of accelerated breeding strategies, such as speed breeding (Hickey et al. 2017; Watson et al. 2018).

### Deflating the bubble of within-family variance

Ever since the introduction of the concept of UC by Schnell and Utz (1975), researchers have been investigating how to exploit it to capture superior outliers from families with high variability. Lehermeier *et al*. (2017) and Allier *et al*. (2019) showed that greater gains can be achieved when mating is based on UC. However, there is also a strong body of evidence in the literature that points to the unexpected and underwhelming result after using the UC (Lado et al. 2017; Wolfe et al. 2021; Bernardo 2022; Wang et al. 2025). Zhong and Jannink (2007) and Wang *et al*. (2025) concluded that UC improved breeding outcomes relative to the mean, only in rare scenarios of high heritability, high selection intensity, and a small number of QTL. In addition, high ratios of 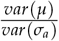 limit the benefit of accounting for within-family variance.

In our study, we found no significant differences between scenarios that accounted for within-family variability and their counterparts in terms of genetic gains, saved diversity, realized inbreeding, and conversion efficiency (Figures 6, 7, and S3). The ratios of 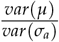 started as high as 299 and 334 in line and hybrid programs and then continued to decline over the years till they reached 61 and 135, which are still too high for within-family variance to make a difference (Figure 8 and Table S4). The initial ratio observed in the line program closely matches the empirical value of 312 reported for wheat yield by Lado *et al*. (2017). We recorded the UI across the years and found that these ratios in our simulations corresponded to negligible additional gains of 0.3–0.5% per cycle on average (Figure 9). In contrast, Lehermeier *et al*. (2017) simulations had much smaller ratios, ranging between 5.3 and 14.25, which explains the additional gain they achieved using the UC.

### Future implications and challenges

This study demonstrates the broad applicability and advantages of integrating optimal recycling with OGM across different breeding systems, species, trait architectures, and genetic effects. However, our simulations were limited to single-trait directional selection. Future work should explore extensions to multitrait selection frameworks using selection indices (Akdemir *et al*. 2019; Endelman 2023). Moreover, we relied exclusively on gBLUP for GS modeling, without assessing alternative genomic prediction methods such as Bayesian regression or reproducing kernel Hilbert space (RKHS)-based models. Furthermore, the impact of varying the stages of GS application and recycling was not investigated in this study and remains an open question. Despite these limitations, we expect that the relative performance of the evaluated strategies would remain robust to such variations.

## Data availability

The code used to generate the results and reproduce the figures will be made publicly available upon publication on GITHUB.

## Author contribution statement

JIyS conceived the study and played a role in coordination, manuscript drafting, and securing funding. SMM conducted all simulations of breeding programs, developed the transition-matrix framework to predict genotypic frequencies across selfing cycles, performed the analyses and visualizations, and conceived and executed the integrated strategy of optimal recycling and genomic mating. JFG led the development of the `MateR` package in collaboration with SMM and JIyS, derived the usefulness importance metric, and contributed to the study design. All authors jointly interpreted the results and co-wrote the manuscript.

## Appendix 1: Building the selfing transition matrix for diploids

Based on the law of total probability, since each parental genotype class is a partition of all possible classes, the overall probability that a progeny ends up being genotype *h* (event *B*_*h*_) is the average over all parental genotype classes *g*. Formally, with *g, h* ∈ {AA, Aa, aa} and events *A*_*g*_ (parental genotype *g*):

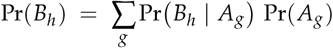

A diploid parent of genotype class *g* carries *K*_*g*_ ∈ {0, 1, 2} copies of allele *A* at a locus; a gamete samples one allele at random. Let *H* ∈ {0, 1} be the number of *A* alleles in the gamete. Then:

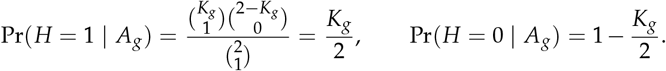

Thus, the gamete allele probabilities are:

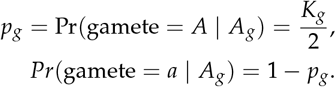

Equivalently in Table 2,

**Table 2.**
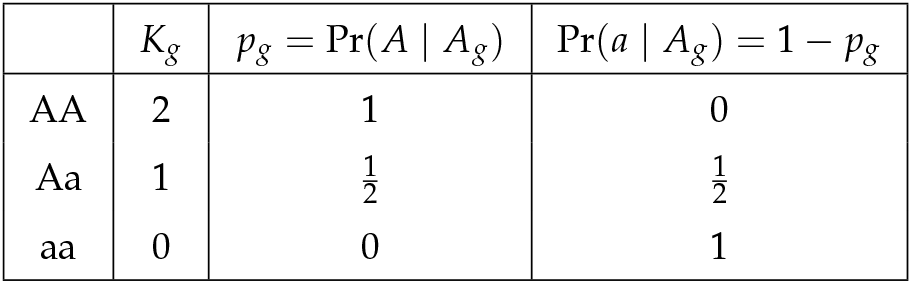
Gamete allele probabilities from a diploid parent in genotype class *g* under random segregation.

Under selfing (random union of two independent gametes from the same parental class *g*), the zygote genotype probabilities are:

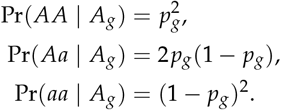

Substituting the values of *p*_*g*_ for each parental class yields the diploid selfing transition matrix *X*(_*ϕ*= 2_) (columns: parental classes *g*; rows: progeny classes *h* in the order *AA, Aa, aa*):

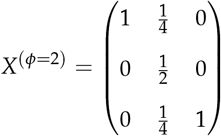

where the columns contain the conditional probabilities of transitioning from the parental to progeny states.

## Supplementary Material

**Figure S1.**
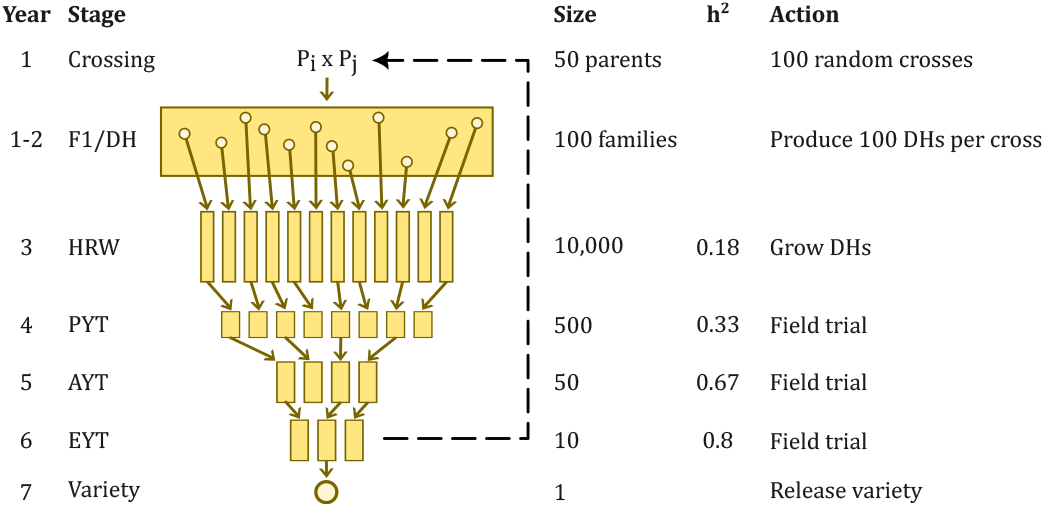
Schematic overview of the line-breeding phenotypic selection (PS) program using double haploid (DH) technology (adapted with modifications from Gaynor *et al*. (2017); Bančič *et al*. (2025)). The program consists of key breeding stages: `DH`, `HRW` (headrows), `PYT` (preliminary yield trial), `AYT` (advanced yield trial), and `EYT` (elite yield trial). Each stage includes varying numbers of individuals (`Size`), yield heritability (*h*^2^), and specific actions. The black dashed line indicates the recycling of individuals from `EYT` into the next breeding cycle.

**Figure S2.**
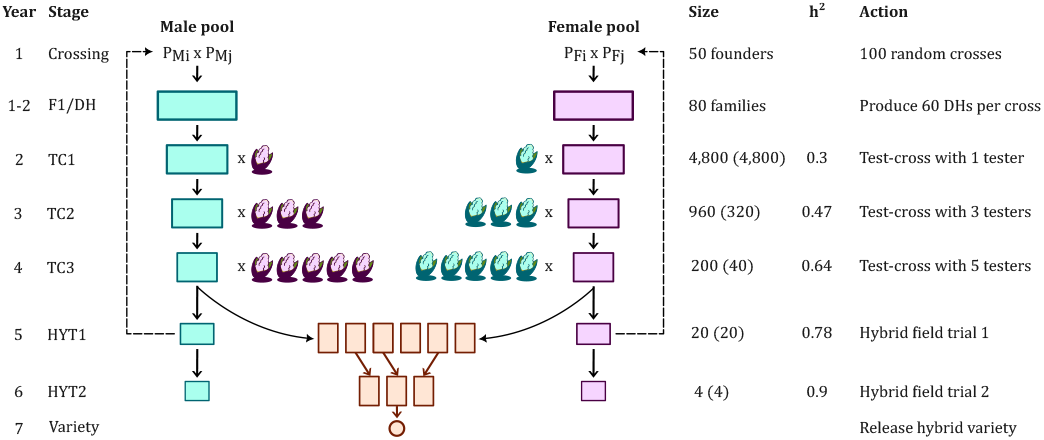
Schematic overview of the hybrid-breeding phenotypic selection (PS) program using double haploid (DH) technology (adapted with modifications from Bernardo (2009); Powell *et al*. (2020); Bančič *et al*. (2025)). The program consists of key breeding stages: `DH`, `TC` (test-cross trial), and `HYT` (hybrid yield trial). Each stage includes varying numbers of individuals (`Size`), yield heritability (*h*^2^), and specific actions. The population size at each `TC` and `HYT` stage refers to the number of hybrids in each heterotic pool, while values in parentheses indicate the number of DH inbreds in each pool. The black dashed line represents the recycling of individuals from `HYT1` into the next breeding cycle.

**Figure S3.**
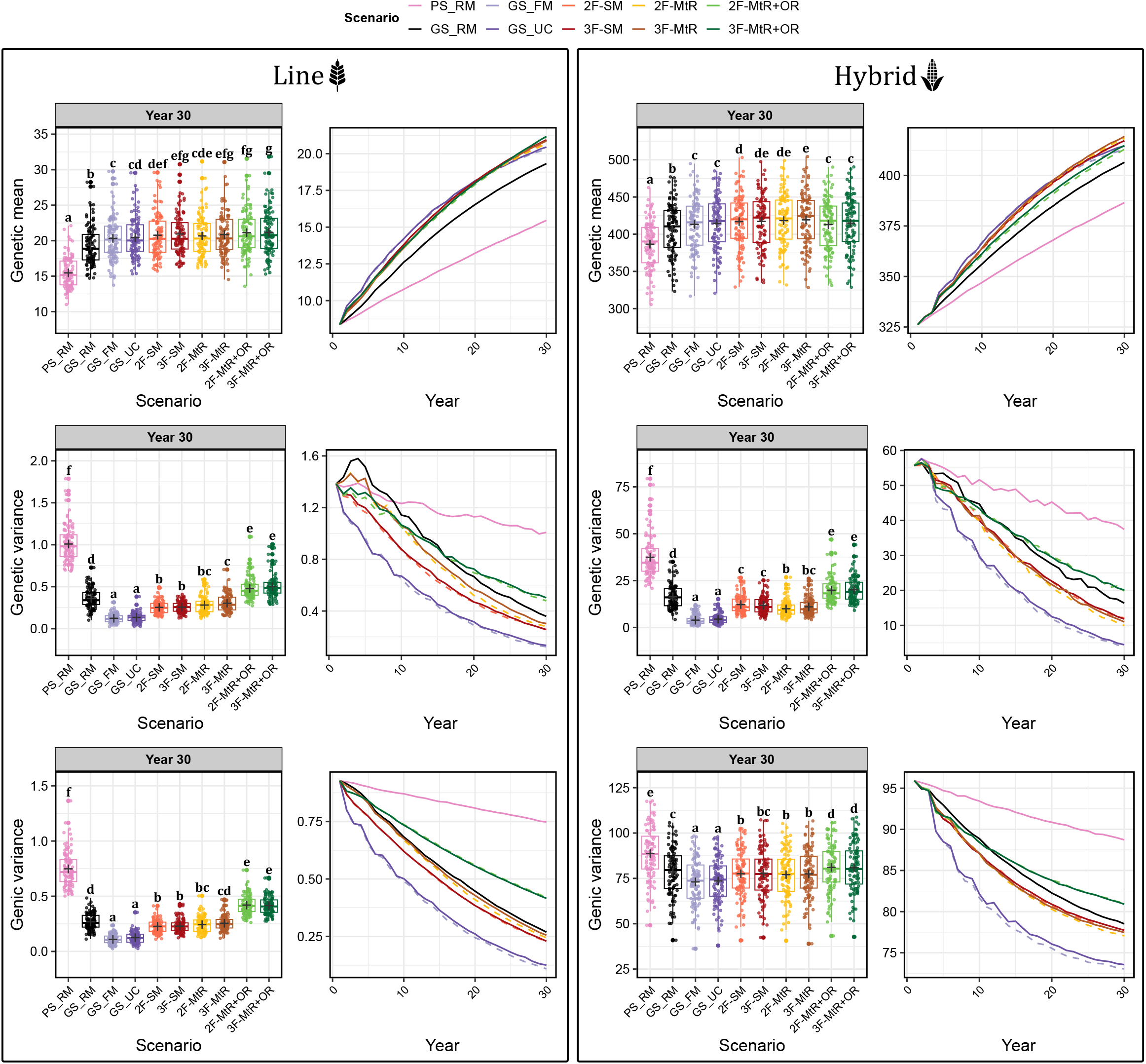
Trends of genetic mean, total genetic variance, and total genic variance for the line and hybrid breeding programs over 30 years of future breeding. Each line represents the mean across 100 simulation replicates. In boxplots, scenarios with the same letter do not differ significantly (Tukey’s HSD test, 99% confidence interval). All scenarios are described in Table 1. Genetic means and variances were recorded at the DH stage for line breeding and test-cross trial 2 (TC2) hybrids for hybrid breeding.

**Table S1.** Average conversion efficiency, achieved standardized gain, and lost genic standard deviation for line and hybrid programs.

| Program | Scenario | Conversion efficiency | Achieved standardized gain | Lost genic standard deviation<sup>a</sup> |
| --- | --- | --- | --- | --- |
| Line | PS_RM | 71.60 ± 10.32 | 7.63 ± 0.66 | 0.098 ± 0.024 |
| Line | 2F-MtR+OR | 39.74 ± 3.55 | 13.51 ± 1.27 | 0.314 ± 0.043 |
| Line | 3F-MtR+OR | 39.36 ± 3.81 | 13.57 ± 1.40 | 0.317 ± 0.043 |
| Line | 3F-MtR | 26.28 ± 3.53 | 13.26 ± 1.28 | 0.458 ± 0.066 |
| Line | 3F-SM | 26.07 ± 3.18 | 13.34 ± 1.11 | 0.484 ± 0.067 |
| Line | 2F-SM | 25.37 ± 3.40 | 13.15 ± 1.27 | 0.486 ± 0.066 |
| Line | 2F-MtR | 25.01 ± 3.38 | 13.05 ± 1.10 | 0.471 ± 0.072 |
| Line | GS_RM | 24.30 ± 3.60 | 11.65 ± 1.06 | 0.445 ± 0.070 |
| Line | GS_UC | 19.46 ± 2.73 | 12.84 ± 1.18 | 0.613 ± 0.082 |
| Line | GS_FM | 18.91 ± 3.27 | 12.69 ± 1.39 | 0.636 ± 0.088 |
| Hybrid | PS_RM | 153.63 ± 43.18 | 6.43 ± 0.90 | 0.391 ± 0.109 |
| Hybrid | 3F-MtR+OR | 115.39 ± 31.94 | 9.32 ± 1.27 | 0.816 ± 0.155 |
| Hybrid | 2F-MtR+OR | 112.91 ± 30.22 | 9.14 ± 1.26 | 0.805 ± 0.169 |
| Hybrid | 3F-SM | 94.89 ± 24.91 | 9.57 ± 1.26 | 0.995 ± 0.174 |
| Hybrid | 2F-SM | 93.28 ± 23.27 | 9.54 ± 1.29 | 1.005 ± 0.179 |
| Hybrid | 3F-MtR | 92.86 ± 24.15 | 9.79 ± 1.27 | 1.010 ± 0.178 |
| Hybrid | 2F-MtR | 89.87 ± 22.23 | 9.68 ± 1.23 | 1.033 ± 0.181 |
| Hybrid | GS_RM | 83.95 ± 18.87 | 8.47 ± 1.12 | 0.950 ± 0.137 |
| Hybrid | GS_UC | 72.70 ± 17.28 | 9.30 ± 1.26 | 1.234 ± 0.195 |
| Hybrid | GS_FM | 68.93 ± 16.18 | 9.15 ± 1.21 | 1.264 ± 0.199 |

<sup>a</sup> Values are reported as mean ± standard deviation across 100 simulation replicates.

**Table S2.** Summary of realized genetic means, gains, genetic variance, and genic variance in the last future breeding year (30) of line and hybrid programs.

| Program | Scenario | Genetic mean | Genetic gain | Total genetic variance | Total genic variance<sup>a</sup> |
| --- | --- | --- | --- | --- | --- |
| Line | PS_RM | 15.46 ± 2.16 | 7.10 ± 1.22 | 1.008 ± 0.217 | 0.748 ± 0.158 |
| Line | GS_RM | 19.32 ± 2.77 | 10.96 ± 1.87 | 0.359 ± 0.118 | 0.269 ± 0.073 |
| Line | GS_FM | 20.33 ± 3.02 | 11.96 ± 2.17 | 0.123 ± 0.051 | 0.109 ± 0.043 |
| Line | GS_UC | 20.45 ± 2.82 | 12.08 ± 1.93 | 0.134 ± 0.056 | 0.125 ± 0.052 |
| Line | 2F-MtR | 20.67 ± 2.95 | 12.30 ± 2.04 | 0.281 ± 0.097 | 0.243 ± 0.072 |
| Line | 2F-SM | 20.77 ± 3.06 | 12.41 ± 2.17 | 0.253 ± 0.076 | 0.227 ± 0.057 |
| Line | 3F-MtR | 20.87 ± 3.00 | 12.50 ± 2.11 | 0.303 ± 0.105 | 0.255 ± 0.067 |
| Line | 3F-SM | 20.94 ± 2.97 | 12.58 ± 2.09 | 0.257 ± 0.065 | 0.229 ± 0.058 |
| Line | 2F-MtR+OR | 21.13 ± 3.13 | 12.76 ± 2.24 | 0.479 ± 0.126 | 0.420 ± 0.086 |
| Line | 3F-MtR+OR | 21.17 ± 3.13 | 12.81 ± 2.26 | 0.502 ± 0.140 | 0.416 ± 0.083 |
| Hybrid | PS_RM | 386.53 ± 32.05 | 60.37 ± 10.59 | 37.44 ± 11.26 | 88.72 ± 14.44 |
| Hybrid | GS_RM | 406.50 ± 34.46 | 80.35 ± 13.35 | 16.36 ± 6.27 | 78.52 ± 13.33 |
| Hybrid | GS_FM | 413.17 ± 35.42 | 87.01 ± 14.45 | 3.90 ± 2.51 | 73.04 ± 12.54 |
| Hybrid | GS_UC | 414.55 ± 35.48 | 88.39 ± 14.59 | 4.47 ± 2.68 | 73.56 ± 12.53 |
| Hybrid | 2F-MtR | 418.34 ± 35.93 | 92.18 ± 14.75 | 9.98 ± 4.06 | 77.06 ± 13.18 |
| Hybrid | 2F-SM | 416.93 ± 36.07 | 90.77 ± 15.20 | 12.22 ± 4.67 | 77.53 ± 13.02 |
| Hybrid | 3F-MtR | 419.31 ± 35.98 | 93.15 ± 14.97 | 10.99 ± 4.43 | 77.47 ± 13.37 |
| Hybrid | 3F-SM | 417.20 ± 35.33 | 91.04 ± 14.56 | 11.83 ± 4.57 | 77.73 ± 13.48 |
| Hybrid | 2F-MtR+OR | 412.95 ± 35.13 | 86.79 ± 14.28 | 19.85 ± 6.68 | 81.08 ± 13.26 |
| Hybrid | 3F-MtR+OR | 414.78 ± 36.09 | 88.62 ± 15.09 | 20.10 ± 6.36 | 80.91 ± 13.57 |

<sup>a</sup> Values are reported as mean ± standard deviation across 100 simulation replicates.

**Table S3.** Average realized inbreeding rates recorded from pedigree (Δ*F*_*A*_) and genomic (Δ*F*_*G*_) information over 30 years of future breeding in line and hybrid programs.

| Program | Parental pool | Scenario | $\Delta F_A(\%)$ | $\Delta F_G(\%)^a$ |
| --- | --- | --- | --- | --- |
| Line | – | PS_RM | $0.319 \pm 0.093$ | $0.594 \pm 0.284$ |
| Line | – | GS_RM | $2.049 \pm 0.365$ | $3.968 \pm 0.915$ |
| Line | – | GS_FM | $3.296 \pm 0.770$ | $6.140 \pm 1.963$ |
| Line | – | GS_UC | $3.077 \pm 0.770$ | $5.620 \pm 1.478$ |
| Line | – | 2F-SM | $2.176 \pm 0.346$ | $4.414 \pm 0.920$ |
| Line | – | 3F-SM | $2.132 \pm 0.368$ | $4.395 \pm 0.974$ |
| Line | – | 2F-MtR | $2.428 \pm 0.476$ | $4.577 \pm 1.054$ |
| Line | – | 3F-MtR | $2.268 \pm 0.459$ | $4.399 \pm 1.041$ |
| Line | – | 2F-MtR+OR | $1.359 \pm 0.261$ | $2.677 \pm 0.604$ |
| Line | – | 3F-MtR+OR | $1.371 \pm 0.239$ | $2.687 \pm 0.592$ |
| Hybrid | Male | PS_RM | $0.462 \pm 0.070$ | $0.888 \pm 0.154$ |
| Hybrid | Male | GS_RM | $1.717 \pm 0.280$ | $3.608 \pm 0.743$ |
| Hybrid | Male | GS_FM | $3.446 \pm 0.821$ | $6.497 \pm 1.777$ |
| Hybrid | Male | GS_UC | $3.106 \pm 0.804$ | $6.100 \pm 1.781$ |
| Hybrid | Male | 2F-SM | $1.937 \pm 0.329$ | $4.192 \pm 0.908$ |
| Hybrid | Male | 3F-SM | $1.912 \pm 0.324$ | $4.266 \pm 0.943$ |
| Hybrid | Male | 2F-MtR | $2.378 \pm 0.470$ | $4.717 \pm 1.110$ |
| Hybrid | Male | 3F-MtR | $2.159 \pm 0.378$ | $4.500 \pm 1.016$ |
| Hybrid | Male | 2F-MtR+OR | $1.281 \pm 0.271$ | $2.607 \pm 0.580$ |
| Hybrid | Male | 3F-MtR+OR | $1.291 \pm 0.273$ | $2.671 \pm 0.608$ |
| Hybrid | Female | PS_RM | $0.465 \pm 0.069$ | $0.896 \pm 0.152$ |
| Hybrid | Female | GS_RM | $1.767 \pm 0.292$ | $3.584 \pm 0.846$ |
| Hybrid | Female | GS_FM | $3.411 \pm 1.001$ | $6.175 \pm 1.648$ |
| Hybrid | Female | GS_UC | $3.259 \pm 0.929$ | $5.902 \pm 1.540$ |
| Hybrid | Female | 2F-SM | $1.920 \pm 0.351$ | $4.135 \pm 0.919$ |
| Hybrid | Female | 3F-SM | $1.902 \pm 0.334$ | $4.059 \pm 0.839$ |
| Hybrid | Female | 2F-MtR | $2.274 \pm 0.501$ | $4.577 \pm 1.024$ |
| Hybrid | Female | 3F-MtR | $2.112 \pm 0.408$ | $4.410 \pm 1.035$ |
| Hybrid | Female | 2F-MtR+OR | $1.297 \pm 0.267$ | $2.635 \pm 0.611$ |
| Hybrid | Female | 3F-MtR+OR | $1.296 \pm 0.280$ | $2.668 \pm 0.632$ |

<sup>a</sup> Values are reported as mean ± standard deviation across 100 simulation replicates.

**Table S4.**
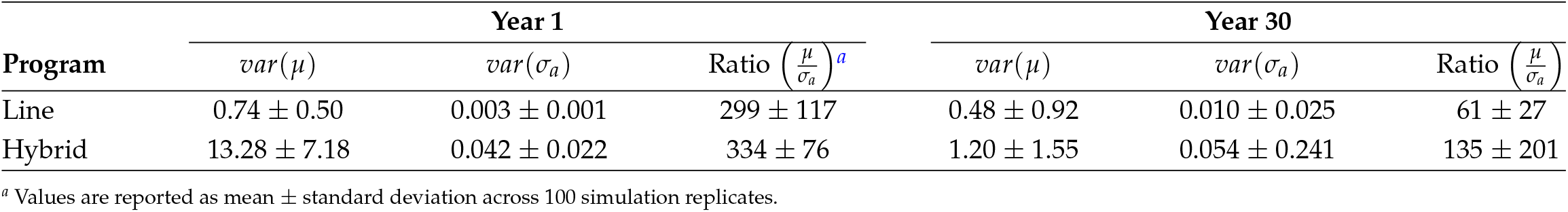
Summary of family genetic mean variance (var(*µ*)), family genetic standard deviation variance (var(*σa*)), and their ratio across the simulated breeding programs at the beginning (Year 1) and end (Year 30) of the future breeding phase.

